# Spatial connectivity for local cortical homogeneity

**DOI:** 10.1101/2025.09.29.679376

**Authors:** Ping Wang, Xi-Nian Zuo

## Abstract

Understanding the functional organization of the primate cortex requires metrics that capture both the temporal and topological dimensions of functional connectivity. Here we propose the spatial connectivity for local homogeneity in cortex (**SoHo**), a vertex-wise, continuous metric that quantifies the degree to which a cortical vertex and its immediate neighbors share similar spatial profiles of whole-brain functional connectivity. We validated **SoHo** using large-scale wakeful resting-state fMRI datasets from the Human Connectome Project (HCP) and the NIH Marmoset Brain Mapping Project. In humans, **SoHo** values showed a striking correspondence with the parcellation boundaries of the HCP multimodal atlas, with low-value regions consistently aligning with areal boundaries. Higher-order association areas exhibited lower **SoHo** values (functional diversity), while primary sensorimotor areas demonstrated higher values (functional uniformity). Cross-species **SoHo** mapping revealed that this primary-to-association gradient is evolutionarily conserved across primates, alongside species-specific adaptations in frontoparietal and motor regions. By capturing the local concordance of spatial fingerprints of whole-brain connectivity, **SoHo** bridges discrete parcellation schemes and continuous models of brain function, offering new insights into primate brain organization and evolution.

## Introduction

Functional parcellation of the human brain refers to the practice of dividing the brain into distinct regions based on specific functional properties, often observed using fMRI or other neuroimaging techniques [1]. Although these functional divisions are widely used and have proven useful in understanding brain organization, there are ongoing debates about whether they are “real” in the sense of being absolute or biologically fixed [2–4]. In the current body of research, functional parcellations can be seen as representative of brain organization rather than rigid universal boundaries [5]. First, brain regions are not necessarily partitioned in identical ways between individuals: functional characteristics can vary according to genetic [6, 7], developmental, and experiential factors (e.g., age, education, or disease) [8, 9], suggesting that functional parcellation is not a hard and fast rule, but a useful approximation. Second, the organization of brain function can change depending on context, task demands, and cognitive state [10]. Resting-state networks can look different from those engaged during a specific task, and even task-based parcellations can change depending on the cognitive function being assessed, implying more flexible than rigidly defined zones. Third, the precision of parcellation methods, whether based on task-/resting-state, or naturalistic fMRI, can vary considerably. Different analytical strategies can lead to different definitions of network regions [11, 12]. The absence of a universally agreed-upon methodology contributes to persisting uncertainty about the true nature of functional boundaries.

These considerations underscore the need for more flexible, continuous representations of brain function that move beyond rigid network boundaries and better capture complex spatiotemporal dynamics and functional organization. A critical step toward this goal has been the development of local functional connectivity metrics. Among these, Regional Homogeneity (ReHo) [13], introduced by Zang and colleagues, represents a seminal contribution: it quantifies the local temporal synchronization of the BOLD signal by computing Kendall’s Coefficient of Concordance (KCC) between the time series of a given voxel and its nearest spatial neighbors. ReHo has since been extensively validated as a reliable, noise-robust and biologically meaningful index of local functional organization in health and disease, and has been systematically reviewed as a multimodal, multiscale neuroimaging marker of the human connectome [14]. Subsequent extensions of this framework, including its implementation on cortical surfaces (2D-ReHo) [15], have demonstrated that local functional homogeneity characterizes hierarchical information processing gradients throughout the cortex, with higher ReHo in primary areas and lower ReHo in higher-order association areas, a gradient consistent with the known complexity of cortical computation [14, 16].

Although informative in describing the homogeneity of temporal profiles within a spatially constrained area, ReHo does not include this area’s entire brain connectivity topology. This connectivity topology has been the driving source of all the parcellation methods of human brain function [1, 17, 18]. Brain connectivity patterns are shaped by two complementary factors. The first is a spatial (geometric) factor, reflecting the tendency of anatomically neighboring regions to share similar connectivity due to wiring cost constraints. The second is a homophily (topological) factor, reflecting the tendency of regions with similar whole-brain connectivity profiles to be preferentially connected [8, 19]. The interplay between these two factors has been consistently demonstrated in species and in the development stages. However, ReHo only captures the spatial factor and measures the concordance of local time series shaped by anatomical proximity. It does not capture whether neighboring vertices share similar whole-brain functional connectivity profiles, that is, whether they are similarly wired to the rest of the brain in a topological sense. This distinction is non-trivial: two neighboring vertices may exhibit highly coherent local time series (high ReHo) while projecting to entirely different large-scale network targets, or conversely, may share nearly identical fingerprints of functional connectivity [20, 21] despite modest local temporal correlation. Bridging this gap requires a metric that integrates the dimension of homophily on whole-brain connectivity into the local neighborhood framework, moving beyond local temporal homogeneity toward local spatial connectivity concordance.

Here, we propose spatial connectivity for local cortical homogeneity (**SoHo**) as a principled extension of the previous ReHo framework that incorporates both spatial and homophily factors of local connectivity. Whereas ReHo computes KCC across the time series of neighboring vertices, SoHo computes KCC across the whole-brain functional connectivity profiles of a vertex and its neighbors. This measures the degree to which immediately adjacent vertices share similar spatial connectivity fingerprints with the rest of the brain. In this sense, **SoHo** inherits the concept of local homogeneity and KCC-based computation from ReHo while extending it from the temporal domain to the spatial domain. **SoHo** captures both the temporal structure of the BOLD signal (inherited through mapping functional connectivity) and the spatial structure of large-scale network organization (captured through the whole-brain functional connectivity profile), constituting a genuinely spacetime concordance metric. Lower **SoHo** values indicate that a vertex and its neighbors have heterogeneous profiles of functional connectivity maps, reflecting functional diversity; higher values reflect a uniform local connectivity fingerprint, indicating functional uniformity. **SoHo** is computed vertex-wise using KCC, a nonlinear, rank-based statistic, and thus is inherently noise-adaptive and robust to outliers, making it suitable for large-scale primate neuroimaging.

We validated this novel approach using large-scale datasets from the Human Connectome Project (HCP) [22] and the NIH Marmoset Brain Mapping Project [23], and addressed the following questions: (1) Does the spatial distribution of **SoHo** values correspond to established parcellation boundaries in the human cortex? (2) Does **SoHo** differentiate higher-order association networks from primary sensorimotor networks in a manner consistent with known functional hierarchies? (3) Are these organizational principles conserved between primate species, and what species-specific adaptations can be identified?

## Results

The spatial distribution of **SoHo** values across the human cortex is presented in Figure 1, overlayed with the refined parcellation boundaries of the HCP 180-region atlas [24]. We observed a striking correspondence between parcellation boundaries and local functional gradient as measured by **SoHo**, with boundary lines consistently aligned with regions that exhibit lower **SoHo**. Consistent with the findings of Gordon et al. [25], our **SoHo**-based analysis successfully delineates the motor cortex into functionally distinct subregions, with boundary lines effectively separating the motor effector and inter-effector regions. However, the visual cortex presents a notable exception, where the **SoHo** metric shows less distinct patterns and the parcellation boundaries do not align as clearly with functional transitions.

**Fig. 1.**
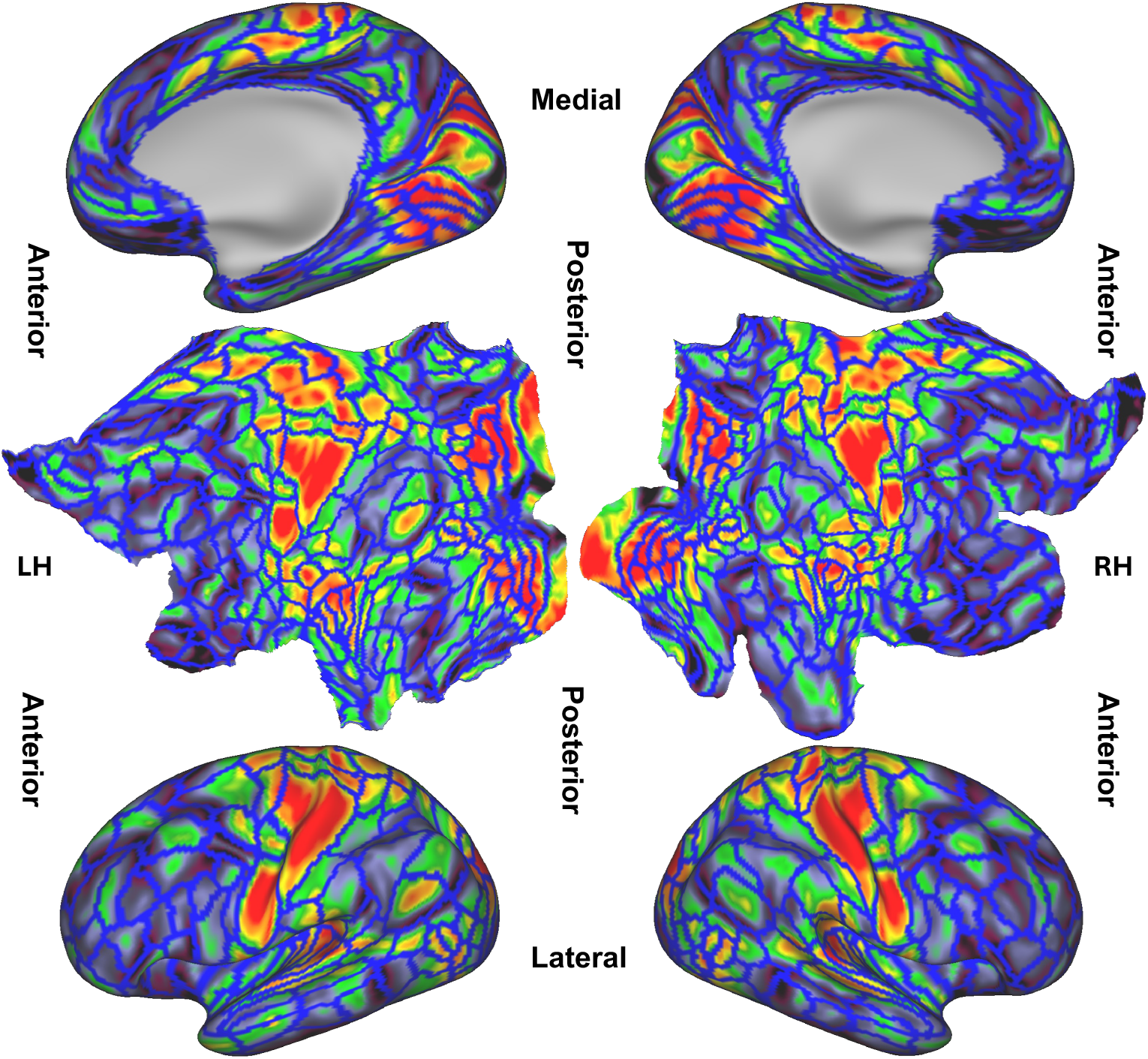
Spatial connectivity mapping for local functional homogeneity (SoHo) in the human cortex. The group-level **SoHo** maps derived from HCP adults are rendered onto the fsLR 32k surface model of both left hemisphere (LH) and right hemisphere (RH). The lateral, medial and flattened cortical surfaces are presented for both hemispheres. Blue lines indicate the boundaries of the HCP-360 multimodal parcellation [24].

To further characterize the functional spacetime concordance in the human cortex or local cortical homogeneity, we extracted mean **SoHo** values from all vertices within each of the 200 parcellation units [18] assigned to 15 canonical cortical networks defined in DU15NET [17] for each hemisphere. These values were plotted along a spiral trajectory in ascending order, as shown in the upper panel of Figure 2. The analysis revealed that regions with lower **SoHo** values (positioned inside the spiral) were predominantly located in higher-order association networks, including the salience and parietal memory network (SAL/PMN), default-mode network (DMN-B), frontoparietal network (FPN-A), and language network (LANG). This pattern indicates a high functional diversity within these networks, consistent with their roles in complex cognitive processes. In contrast, primary sensorimotor and sensory networks, including somatomotor networks (SMOT-A and SMOT-B) and visual peripheral networks (VIS-P), displayed higher **SoHo** values (positioned outside the spiral), reflecting the functional uniformity characteristic of these fundamental processing areas. A particularly important observation emerged within the motor cortex, where regions sharing the same network assignment (indicated by identical color coding) displayed a wide range of **SoHo** values. Some regions exhibited low **SoHo** values that indicate high functional diversity, while others demonstrated high **SoHo** values that reflect functional uniformity.

**Fig. 2.**
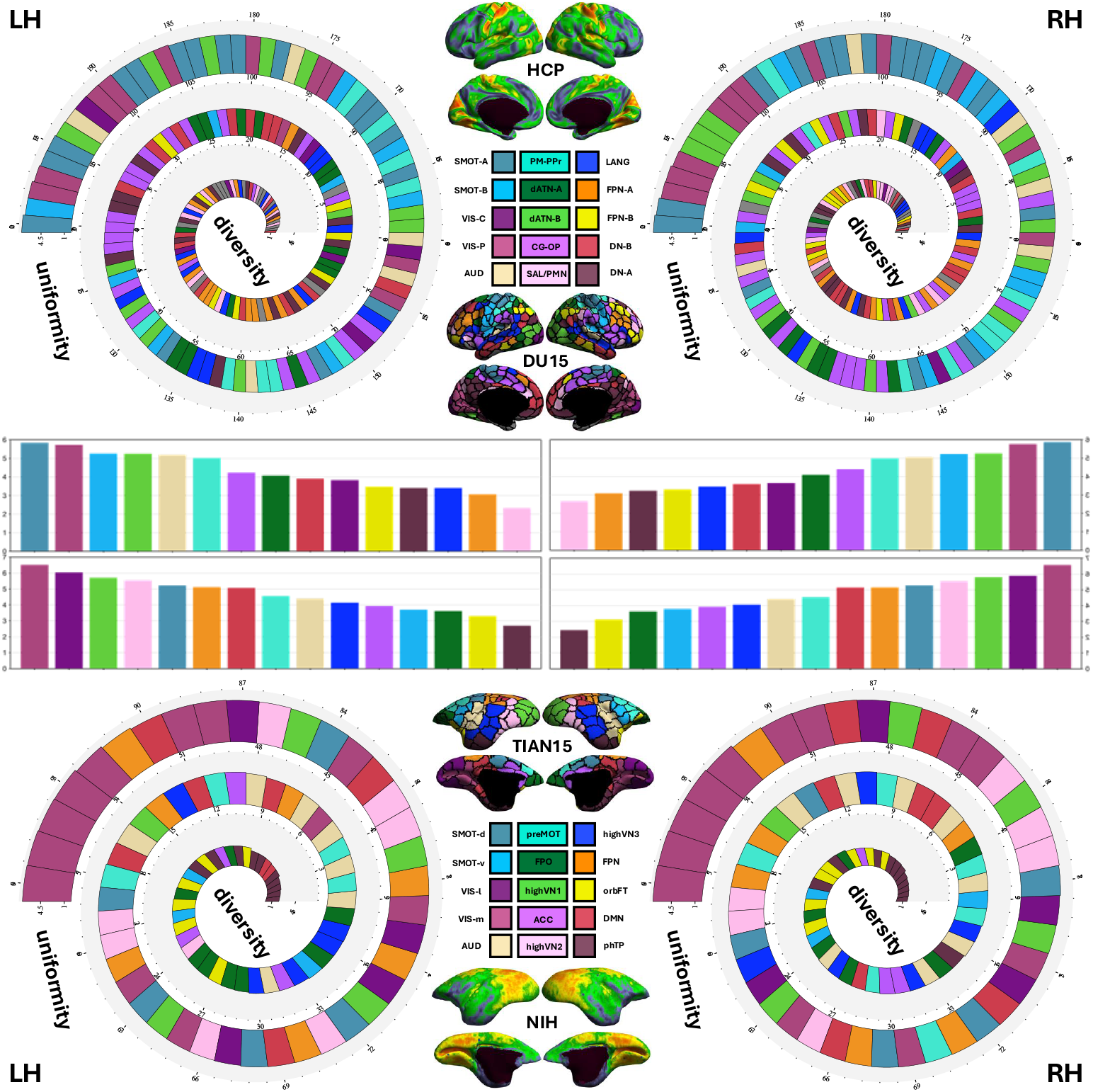
SoHo variation in the primate cortex. An organization of local function from uniformity to diversity is depicted across large-scale networks and parcellations of primate cortex (top: human, bottom: marmoset). Details of the SoHo mapping and its spiral ordering are documented in the main text for human and marmoset, respectively.

The **SoHo** analysis was extended to the marmoset cortex using a similar approach, extracting mean **SoHo** values from all vertices within each of the 96 parcellation units assigned to fifteen canonical cortical networks defined by Tian et al. (2022) [26]. As shown in the lower panel of Figure 2, in marmosets, regions with lower SoHo values (inside the spiral) were located primarily in the parahippocampus and temporal pole network (phTP), frontal pole network (FPO) and orbital frontal network (orbFT). In particular, high-level visual network 3 (High-VN3), which corresponds to areas related to language in humans, also showed relatively low **SoHo** values. Higher **SoHo** values were predominantly found in networks associated with primary cortical areas, particularly lateral and medial visual networks (VIS-l and VIS-m). The frontoparietal-like network (FPN) in marmosets also showed relatively high SoHo values, suggesting a more uniform functional organization compared to the human FPN.

## Discussion

Functional parcellations represent pragmatic representations of how the brain operates at the system level, providing valuable information for understanding brain function [1]. Although they are real in the sense that they capture meaningful organizational principles, they are not absolute fixed regions. Brain organization is likely more fluid, with regions and networks dynamically interacting, and parcellations reflecting approximations rather than strict anatomical or functional boundaries [20]. However, current brain parcellation methods typically regard brain parcellation as rigid and discrete, often overlooking the continuous nature of functional transitions across the cortex [12, 27–29]. In this study, we propose **SoHo**, a vertex-wise continuous metric of cortical functional organization. Building on ReHo [13], which characterizes local connectivity through the temporal synchrony of BOLD time series, **SoHo** extends this approach into the homophily dimension. Rather than measuring only whether neighboring vertices fluctuate synchronously over time, **SoHo** quantifies whether they share similar whole-brain connectivity patterns. This enables **SoHo** to capture spatiotemporal brain concordance, providing a more precise and dynamic characterization of functional parcellations.

Our analysis demonstrated a striking correspondence between the parcellation boundaries of the HCP atlas and regions of low **SoHo** values, indicating that functional boundaries naturally emerge at locations where regional functional uniformity transitions to diversity. This pattern validates **SoHo** as an effective metric for characterizing functional organization and accurately mapping complex functional architecture. In humans, higher-order association networks exhibited lower **SoHo** values indicating functional diversity, while primary sensorimotor networks displayed higher **SoHo** values reflecting uniformity. This gradient from primary to association cortex is consistent with established hierarchical models of cortical organization [16, 30], and echoes the pattern previously reported for ReHo [14]. Critically, **SoHo** extends these findings by showing that the primary-to-association gradient is reflected not only in local temporal coherence but also in the local concordance of whole-brain connectivity fingerprints.

In contrast to the correspondence observed across the rest of the cortex, the visual cortex presents a notable exception. Although **SoHo** shows less distinct patterns in this region, the existing low-value areas do not align clearly with the HCP parcellation boundaries. This inconsistency is most parsimoniously attributed to the multimodal nature of the HCP parcellation scheme itself, which integrates not only functional connectivity, but also structural features including cortical thickness, myelination patterns (T1w/T2w ratio), cortical surface area and task activation maps [31]. This suggests that the HCP atlas is not a pure functional brain parcellation, but rather a composite of various brain modalities. Multiple neuroanatomical measures (functional connectivity, cortical thickness, myelination) may provide redundant information, limiting the unique contribution of functional measures.

Comparative analysis between human and marmoset cortex reveals both evolutionary conservation and species-specific adaptations in functional organization. The general principle that primary sensory areas exhibit greater functional uniformity while association areas display greater diversity appears to be conserved between primate species, suggesting fundamental organizational constraints in mammalian brain evolution [32]. This conservation likely reflects selective pressure for efficient sensory processing that is critical for survival across species. However, important species differences emerged that reflect distinct evolutionary pressures and cognitive specializations. The relatively high **SoHo** values in marmoset FPN contrast with the functional diversity observed in human FPN, potentially reflecting the expanded executive control capabilities that characterize human cognition [33]. This difference may reflect the evolutionary expansion and increased complexity of executive control networks in humans. In contrast, the greater functional diversity in the marmoset motor regions may reflect the increased demands for motor control of the marmoset locomotion and arboreal lifestyle [34]. The specialized motor requirements for navigating complex three-dimensional environments necessitate more functionally diverse motor control networks compared to human bipedal locomotion. High-VN3, the regions corresponding to areas of human language [35], showed different functional characteristics in marmosets. These regions, responsible for face recognition, vocalization processing, and social cognition in marmosets [36, 37], exhibited low **SoHo** indicating functional diversity. This finding provides valuable information on the evolutionary trajectory of language-related brain regions and their precursor functions in nonhuman primates.

Although assigned to a single functional network, it is heterogeneity within the motor cortex. The wide range of **SoHo** values within the motor cortex reflects the varied computational demands of different motor functions, from simple reflexive movements to complex skilled behaviors requiring extensive cortical coordination [38, 39]. This variability challenges the traditional view of the motor cortex as a homogeneous functional unit and supports more nuanced models of motor control that incorporate multiple specialized subregions. This finding suggests that current parcellation schemes may oversimplify the functional organization of motor regions and supports the need for more continuous spacetime concordance representations of brain function rather than discrete boundaries. The successful delineation of motor effector and inter-effector regions [25] using **SoHo** demonstrates the utility of **SoHo** to identify functionally meaningful subdivisions within traditionally defined networks.

In summary, the **SoHo** metric provides a novel lens for understanding brain functional organization that bridges the gap between discrete parcellation schemes and continuous models of brain function. By revealing the spacetime concordance of functional coherence across the primate cortex, **SoHo** offers new insights into the evolution of brain networks. Our findings support a more nuanced view of brain organization that emphasizes functional transitions rather than rigid boundaries, with important implications for both basic neuroscience and clinical applications. Successful cross-species validation of **SoHo** in humans and marmosets establishes its utility for translational research and provides a foundation for future comparative studies of primate brain evolution and function. This framework represents a critical step toward reconciling the apparent contradiction between parcellation schemes and the continuous nature of brain organization, offering new tools to understand healthy or disease brain function.

## Methods

### Participants and datasets

Wakeful resting-state fMRI (rfMRI) data were obtained from two large-scale open resources: the HCP (N = 1,003) [22] and the NIH Marmoset Brain Mapping Project (N = 26) [23]. The HCP dataset consists of healthy young adults (ages 22–35) scanned at Washington University in St. Louis on a customized 3T Siemens Connectome Skyra scanner. Each participant completed two rfMRI sessions on separate days, with two runs per session (one left-to-right and one right-to-left phase-encoding direction), yielding four runs of approximately 14.4 minutes each (TR = 720 ms, TE = 33.1 ms, 72 slices, 2 mm isotropic voxels, 1,200 volumes per run). The marmoset dataset was acquired as part of the NIH Marmoset Brain Mapping Project [23], which provides fully awake, head-fixed rfMRI data for common marmosets (Callithrix jacchus). All marmoset data were collected using a 9.4T Bruker BioSpec scanner (TR = 2,000 ms, TE = 25 ms, 0.5 mm isotropic voxels). All data used in this study were obtained from publicly available repositories and are fully de-identified; no additional ethical approvals were required.

### Data preprocessing

For the HCP dataset, minimally preprocessed data in the CIFTI space (91,282 vertices on a standard 32k surface mesh) were used as provided by the HCP consortium [31]. The HCP minimal preprocessing pipeline includes gradient-nonlinearity correction, motion correction using FSL’s MCFLIRT, EPI distortion correction using fieldmaps, registration to the MNI152 template, brain-boundary-based registration of functional to structural data, and surface registration using FreeSurfer and MSMAll [31]. Nuisance signal regression employed ICA-FIX denoising to remove structured artifacts [40], followed by removal of the mean time series from white matter and cerebrospinal fluid. No spatial smoothing was applied prior to connectivity computation. A high-pass temporal filter with a cutoff of 2,000 s was applied to remove slow signal drifts.

For the marmoset dataset, the preprocessing was done by using the Marmoset Brain Mapping pipeline [23, 26], which includes motion correction, EPI distortion correction, co-registration of individual anatomical images and nonlinear registration in the NIH Marmoset Brain Atlas. Nuisance regression included removal of motion parameters (six rigid-body parameters plus their first derivatives), white matter, and cerebrospinal fluid signal. The preprocessed data were projected onto a standard marmoset cortical surface with 76,457 vertices.

All data used in this study were obtained from publicly available repositories and had undergone rigorous quality control procedures as part of their respective data collection protocols [22, 23]. No additional motion-based exclusion criteria were applied beyond those implemented in the original preprocessing pipelines.

### Group-level connectome construction

To efficiently aggregate individual rfMRI data into a group-level representation while reducing the influence of random noise, we employed MELODIC’s Incremental Group-Principal Component Analysis (MIGP) [41]. This method concatenates the preprocessed time series from all participants in an incremental, memory-efficient manner and extracts the leading principal components that capture the dominant patterns of shared functional variance. The output is a set of spatial eigenvectors, weighted and re-normalized, that represent the group-level structure of functional covariance. Compared to simple cross-subject averaging of functional connectivity matrices, MIGP better preserves the signal-to-noise characteristics of each individual’s data while capturing group-representative connectivity patterns. The top principal components were retained for HCP (N = 4,500) and for the marmoset dataset (N = 1,800). The output of spatial eigenvectors were then re-normalized, reweighted, and used to compute a group-level dense functional connectivity matrix, resulting in a 91,282 × 91,282 connectivity matrix for humans and a 76,457 × 76,457 matrix for marmosets.

### Functional connectivity profile computation

For each cortical vertex, its functional connectivity profile throughout the brain, also known as its functional “fingerprint” [20], was calculated from MIGP-derived group-level time series. Specifically, the Pearson’s correlation coefficient was calculated between the time series of each target vertex and those of all other cortical and subcortical vertices across the whole brain. The resulting correlation values were then transformed using Fisher’s r-to-z transformation to normalize their distribution and stabilize variance across the range of correlation values, producing the functional connectivity vector of z-values with a length equal to the total number of vertices for each target vertex. These vertex-wise functional connectivity profiles capture the full spatial pattern of functional interactions between each cortical location and the rest of the brain, encoding both the strength and sign (synchrony and asynchrony) of functional relationships.

### SoHo computation

Based on these full-brain functional connectivity profiles, the **SoHo** metric quantifies the degree of spatial connectivity to a given cortical vertex, from its immediate spatial neighbors, or sharing similar functional connectivity profiles. For each target vertex, its nearest neighboring vertices on the cortical surface mesh are identified based on geodesic adjacency. Two neighborhood sizes were typically computed: step-1 neighbors (6 adjacent vertices) and step-2 neighbors (18 adjacent vertices) [14]. Kendall’s Coefficient of Concordance (KCC) *W* [42] of the 7 or 19 functional connectivity profiles of the full-brain was used to measure spatial connectivity for local cortical homogeneity within the area centered on the given vertex (Fig. 3).

**Fig. 3.**
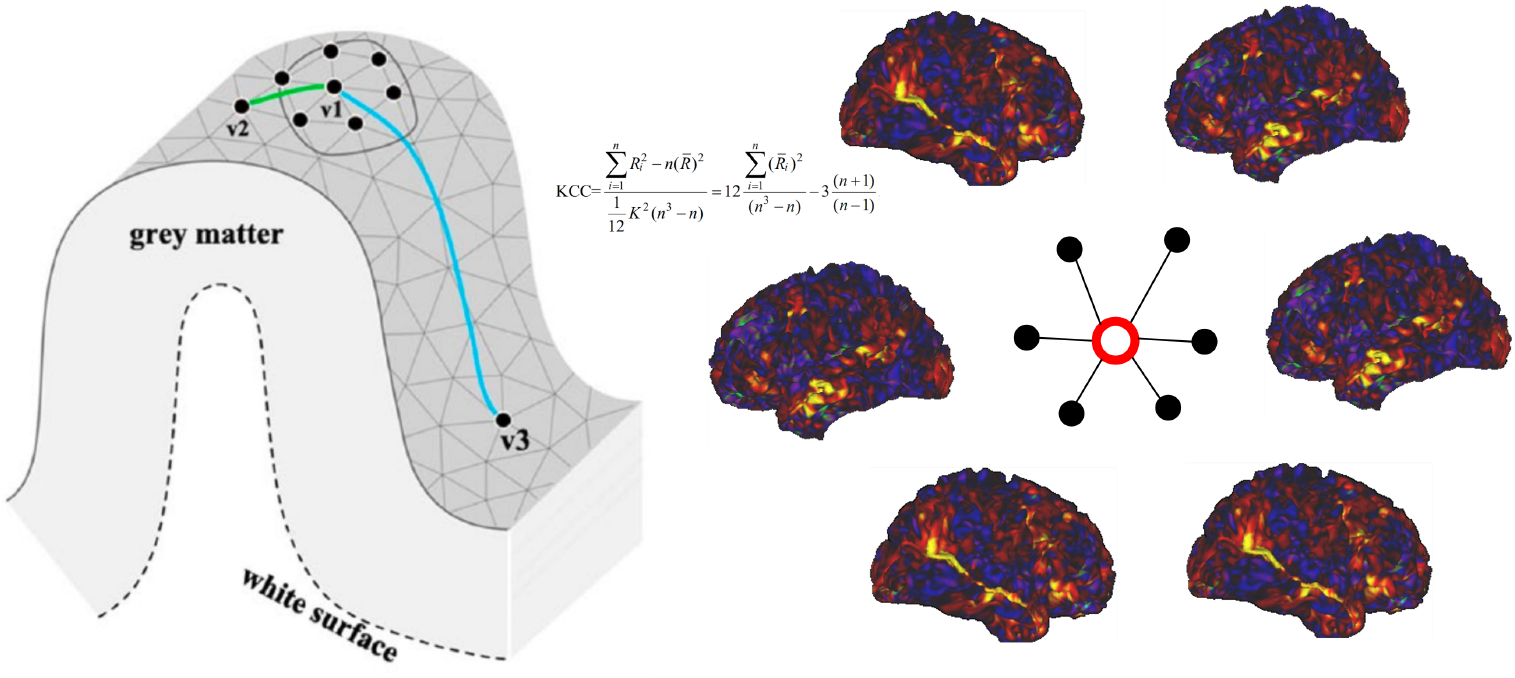
Schematic diagram of the SoHo calculation. Given a vertex v1 on the cortical surface (the left panel), **SoHo** measures the KCC among functional connectivity maps of the seven vertices including v1 (red color) and its six neighboring vertices (black color, the right panel).

The mathematical formula for KCC or *W* is detailed as Equation (1), where *K* is the number of vertices in the neighborhood that includes the target vertex itself (step-1: *K* = 7, which produces **SoHo-1**; step-2: *K* = 19, which produces **SoHo-2**); *n* is the total number of vertices that make up each vertex’s functional connectivity profile; *R*_*i*_ is the sum of the ranks of the z-values at the *i*-th vertex across the neighboring vertices; 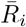 is the mean rank at the vertex *i* between neighbors; and 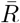 the overall mean rank across all neighboring vertices of *K* and all *n* vertices of the functional connectivity fingerprint. The final **SoHo** was calculated as in Equation (2) to achieve more normally distributed values across the cortex.

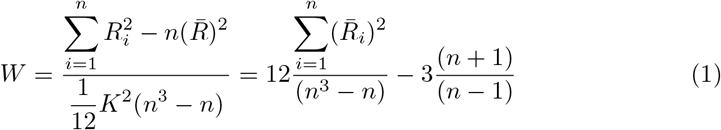

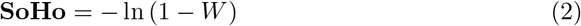

The KCC ranges from 0 to 1, where 0 indicates no concordance among functional connectivity profiles (**SoHo** = 0, minimal spatial connectivity or maximum functional diversity) and 1 indicates perfect concordance (**SoHo** = infinity, maximum spatial connectivity or functional uniformity). Because KCC is rank-based and nonlinear, it is inherently robust to non-normal distributions of connectivity (z-values) and resistant to the influence of outliers, properties that are advantageous for large-scale neuroimaging data [14]. In the present study, **SoHo** was calculated using the step-1 neighborhood (*K* = 7). This choice follows established practice in surface-based local functional homogeneity analysis [13, 14]: a step-1 neighborhood provides the most spatially specific estimate of local functional connectivity concordance, while a larger step-2 neighborhood risks mixing signals from functionally distinct regions across parcellation boundaries, which would precisely attenuate the functional transitions that **SoHo** is designed to capture. All computations were implemented in MATLAB within the Connectome Computation System (CCS) [43].

## Acknowledgments

The authors thank Dr. Cirong Liu from the Institute of Neuroscience, Chinese Academy of Sciences for valuable discussions during the early preparation of the manuscript and sharing the marmoset dataset, and the National Basic Science Data Center for informatic support.

## Declarations

### Author Contributions

P.W. and X.N.Z. designed and performed the experiments, analyzed the data and wrote and edited the manuscript. X.N.Z. developed methods for data processing.

### Data availability

The wakeful rfMRI human data are available from the database of the Human Connectome Project (HCP) (https://db.humanconnectome.org). which is supported by the NIH Blueprint for Neuroscience Research 1U54MH091657 (principal investigators: David Van Essen and Kamil Ugurbil) and the McDonnell Center for Systems Neuroscience at Washington University. The wakeful rfMRI marmoset data are available from the Chinese Color Nest Data Community (CCNDC: https://ccndc.scidb.cn/en) at Science Data Bank (https://doi.org/10.57760/sciencedb.07943 or https://cstr.cn/31253.11.sciencedb.07943).

### Code availability

All codes developed for SoHo are available as part of the Connectome Computation System (CCS: https://github.com/zuoxinian/CCS) via GitHub.

